# Discover Potential New Epitopes through Post-Translational Modification in Sjögren’s Disease

**DOI:** 10.64898/2025.12.30.697079

**Authors:** Danmeng Li, Alexandria Voigt, Cuong Nguyen

## Abstract

Sjögren’s syndrome (SjD) is a chronic autoimmune disorder in which the immune system attacks the glands that produce tears and saliva, leading to symptoms such as dry eyes and dry mouth. If left untreated, SjD can also cause inflammation and damage to other parts of the body, including the skin, lungs, kidneys, and nervous system, and increase the risk of developing lymphoma. The human leukocyte antigen (HLA) class II molecule HLA-DR3 is strongly associated with SjD. To investigate how post-translational modifications (PTMs) influence the presentation of SjD-associated autoantigens by HLA-DR3, we employed a computational framework to analyze the binding of PTM-mimic peptides to HLA-DR3. Our analysis revealed that PTM-mimic substitutions at canonical anchor positions rarely improved predicted binding affinity using the Stabilized Matrix Method, with most modifications resulting in reduced affinity. However, a comprehensive analysis of full-length SjD-associated autoantigen sequences (Ro60, Ro52, La) identified discrete regions with high densities of PTM-eligible anchor sites, specifically, the Ro60 HEAT solenoid, Ro52 RING/B-box/PRY-SPRY modules, and the La motif-RRM1 region, suggesting that PTMs may alter epitope presentation in a sequence-dependent manner. Experimental validation of selected PTM-mimic peptides demonstrated enhanced T cell responses, which were associated with increased binding affinity to HLA-DR3. Structural modeling of a representative complex revealed that PTM-mimic peptides adopt a slightly shifted backbone orientation and altered side-chain positioning, leading to a larger peptide-DR3 interaction interface. These findings provide new insights into the role of PTMs in shaping the immunogenicity of SjD-associated autoantigens and highlight the potential for PTM-mimic peptides to modulate T cell responses in SjD.

## INTRODUCTION

Sjögren’s disease (SjD) is a chronic autoimmune disease characterized by lymphocytic infiltration of the salivary and lacrimal glands, resulting in xerostomia and xerophthalmia, with additional systemic manifestations in a substantial proportion of patients. It is one of the most common connective tissue diseases, affecting approximately 0.5-1% of adults and showing a strong female predominance^1^. Serological detection of autoantibodies against Ro/SSA (Ro60 and Ro52/TRIM21) and La/SSB remains central to diagnosis and is incorporated in the 2016 ACR-EULAR classification criteria^2^. Despite their diagnostic utility, these ribonucleoproteins are ubiquitous intracellular molecules, and the processes that make them immunogenic in SjD remain poorly understood.

Post-translational modifications (PTMs) have emerged as important contributors to immune recognition in several autoimmune diseases. In rheumatoid arthritis, the citrullination of joint proteins generates neoepitopes that stimulate the production of anti-citrullinated protein antibodies^3^. Additionally, carbamylation produces homocitrulline-containing antigens associated with severe phenotypes, even in patients lacking anti-citrullinated protein antibodies (ACPAs)^4^. In systemic lupus erythematosus (SLE), the apoptosis-associated phosphorylation of histones creates immunogenic epitopes^5^, and oxidative modifications, such as 4-hydroxynonenal-modified Ro60, accelerate autoimmunity in vivo^6^. Recent reviews have highlighted that citrullination, acetylation, oxidation, and phosphorylation can reshape self-proteins in ways that promote their recognition by autoreactive lymphocytes^7^. Together, these findings suggest that PTMs can modify the biochemical and immunological properties of self-antigens, thereby contributing to the breakdown of tolerance.

In SjD, several observations suggest a similar role for PTMs in influencing the antigenicity of Ro and La proteins. Ro52 undergoes phosphorylation that affects its E3 ubiquitin ligase activity and may expose regions of the molecule to immune recognition^8^. Oxidized Ro60 promotes accelerated autoimmune responses in experimental systems^9^. More recently, studies in salivary gland epithelial cells (SGECs) have identified dysregulated phosphorylation pathways, including increased ETS1 and phospho-ETS1 levels^10^, and elevated lipid-reactive oxygen species (ROS)-associated STAT4 phosphorylation driven by reduced GPX4 expression^11^. These findings support the idea that Ro60, Ro52, and La are exposed to PTM-rich environments within disease-relevant tissues. However, existing studies largely focus on individual residues or specific signaling pathways and do not define how PTMs distributed along the full length of these antigens might influence the repertoire of peptides available for HLA-DR3 presentation.

Therefore, a broader analytical framework is needed to examine the impact of PTMs across Ro60, Ro52, and La. The key question is whether these modifications reshape the landscape of peptides capable of enhancing antigen presentation by MHC/HLA, specifically SjD-risk alleles, for instance, HLA-DR3. To address this gap, we undertook a comprehensive, sequence-wide in silico analysis comparing predicted HLA-DR3 binding between native and PTM-modified forms, complemented by experimental validation of selected epitopes. This integrated approach provides the foundation for determining whether PTM-driven changes have the potential to influence antigen processing and T-cell response in the autoimmune process of SjD.

## METHODS

### Selection of SjD-associated autoantigenic peptides

Full-length human Ro60, Ro52, and La sequences (Ro60: UniProt P10155, 525 amino acids (aa); Ro52: UniProt P19474, 475 aa; La: UniProt P05455, 408 aa) were scanned using a sliding-window approach that generated overlapping 15-mer peptides at a step size of one residue. This approach follows established computational epitope scanning strategies applied to autoimmune antigens (Backert & Kohlbacher, 2015). The overlapping peptide dataset provided a sequence-wide and unbiased survey of potential HLA-DR3-restricted epitopes of Ro52, Ro60, and La antigens. We also utilized a second peptide dataset using the Stabilized Matrix Method (SMM), which had been previously performed^12^. The SMM-peptide dataset consists of 15-mer peptides computationally predicted to bind HLA-DR3, derived from Ro60, Ro52, and La.

### Binding core prediction and anchor position assignment

For each peptide in overlapping- and SMM-peptide datasets, the predicted 9-mer binding core and the corresponding anchor positions were obtained using the NetMHCIIpan 4.1 binding core predictor^13^ restricted to HLA-DR3. The predicted core was treated as the reference register for subsequent substitution analysis. Anchor residues at positions P1, P4, P6, and P9 identified in this manner were recorded to determine PTM-eligible substitution sites in this study.

### Post-translational modification-mimic substitutions

PTM-mimic residues were introduced only at canonical HLA-DR3 anchor positions and only when a substitution has established biochemical precedent for modeling the charge or polarity associated with the corresponding modification. Deamidation was approximated by replacing glutamine with glutamate, which removes the amide group while preserving side-chain geometry^14^. Citrullination was modeled by substituting arginine with glutamine, a widely used approach to neutralize the guanidinium group and mimic the loss of positive charge^15,16^. Phosphorylation was modeled by replacing serine or threonine with aspartate to introduce a negative charge, a widely used phospho-mimetic strategy^17,18^. Acetylation of lysine was mimicked by substituting lysine with glutamine to eliminate the ε-amino charge while preserving side-chain size^19,20^ (Megee et al., 1990; Wang & Hayes, 2008). Methionine oxidation was approximated by replacing methionine with glutamine, which increases polarity relative to the thioether side chain. This has been used as an oxidation-like mimic in peptide-MHC studies^14^. Aromatic hydroxylation was represented by substituting phenylalanine with tyrosine, which introduces a hydroxyl group while maintaining aromaticity^21^. These six substitution classes comprised the PTM-mimic variants evaluated in this study.

In the SMM-peptide dataset, for each eligible anchor position, a PTM-mimic 15-mer was generated by replacing only the anchor residue while maintaining all other native positions. Each PTM-modified peptide was paired with its corresponding native peptide of identical length and predicted register.

### HLA-DR3 binding prediction

All native and PTM-mimic peptides were evaluated using the IEDB NetMHCIIpan 4.1 BA predictor^13^ restricted to HLA-DRB1*03:01. Predictions were recorded as IC_50_ (nM) and percentile rank. The log₂(Native/PTM IC_50_) values were calculated to quantify both the magnitude and direction of PTM-associated affinity changes. This transformation places increases and decreases in predicted binding on a common fold-change scale, allowing for direct comparison of affinity shifts across peptides with different baseline IC_50_ values. The use of log_2_-transformed fold-change values follows standard practice for comparing peptide-MHC affinity shifts^22^.

### Dataset integration

The native and PTM pairs were analyzed within the sliding-window dataset derived from full-length Ro60, Ro52, and La. The SMM-peptide dataset was mapped to its corresponding positions within the dataset and annotated accordingly. All unique native and PTM pairs were included in the computational analysis pipeline. Structural modeling and in vitro validation were performed on selected peptide pairs.

### Data visualization

Data visualizations for computational analyses were generated using Python 3.10^23^. Dataset handling was performed using Pandas^24^, and plots were created with Matplotlib v3.7^25^. The same axis scaling, smoothing parameters, and color mapping were applied across Ro60, Ro52, and La to ensure comparability among figures.

### Validation of the selected PMT-mimics

Four pairs of native and PTM 15-mer peptides were selected and synthesized (Alan Scientific, MD, USA) at > 90% purity (HPLC-confirmed) and reconstituted at 1 mg/mL in sterile buffer. Screening of PTM peptides was conducted by using splenocytes of NOD.DR3 mice, a congenic strain bred to carry the human DR3 HLA haplotype. Briefly, NOD.DR3 spleens were mashed through a 70 μm strainer, and red blood cells were lysed from the single cell suspension using lysis buffer [0.802% NH4Cl, 0.084% NaHCO3, and 0.037% EDTA] for 13.5 minutes. Each peptide was incubated at 100 ug/ml with 2 × 10^5^ splenocytes (37°C, 5% CO_2_, 48 hours). Where indicated, plates were coated (4 °C, overnight) with 50 µl of 10 µg/ml anti-CD3 (BD Pharmingen, Franklin Lakes, New Jersey). After washing the plate, 5 µg/ml anti-CD28 (BD Pharmingen, Franklin Lakes, NJ) was added prior to other well contents. The supernatant was then collected to measure the secreted IL-2 (BioLegend, San Diego, CA) according to the manufacturer’s instructions. Plate was run on Tecan NanoQuant and results were analyzed with Python 3.

### Determining the specificity of the PTM-mimics using SjD-associated T cell receptor

Lentivirus expressing T cell receptor (TCR) from SjD pathogenic T cells was produced as described previously^1^. To isolate antigen-presenting cells (APC), splenocytes of NOD.DR3 were isolated as described previously, and stained with 3 uL PE anti-CD3, followed by incubation with anti-PE microbeads (Miltenyi Biotec, Germany). Labeled cells were run on a MACS MS column for isolation of APC. Isolated APCs were treated with Mitomycin C (20 min, 37 °C), then plated on a 24-well plate at a density of 2.5 × 10^5^ cells per well. Peptides were added at a concentration of 100 µg/ml. As a negative control for peptide specificity, an unrelated peptide, ribonucleotide reductase subunit 1 from human alphaherpesvirus 1, positions 709 to 721 (NVTWTLFDRDTSM), was utilized. Plates were incubated (5% CO_2_, 37 °C, 48 hours), supplemented with additional media, and then incubated for an additional 24 hours. Flow cytometry was performed using a Cytek Aurora (Cytek Biosciences, Fremont, California) to detect the presence of TCRs via GFP fluorescence. Results were analyzed with FlowJo v10 and graphed on Python 3.10.

### Structural modeling and interface analysis

Structural comparison of native and PTM-mimic peptides was performed for one representative Ro60-derived peptide pair. Predicted HLA-DR3 or HLA-DRB1*03:01-peptide complexes were generated using TCRmodel^26^, which provides template-based modeling of class II peptide-MHC interactions. Models for Native and PTM-mimic peptides were obtained separately and aligned in PyMOL (The PyMOL Molecular Graphics System, Version 2.5 Schrödinger, LLC.) using the DR3 α/β heterodimer as the structural reference. Peptide-HLA interface properties were quantified using PDBsum (RRID:SCR_006511)^27^. PDBsum-generated metrics were extracted directly from the server output without manual modification. Structural figures were prepared in PyMOL.

### Statistical analysis

All quantitative data are presented as mean±SEM. Paired comparisons between native and PTM-modified peptides were analyzed using two-tailed paired t test. Statistical analyses were performed using GraphPad Prism version 10.0.0 for Mac OS X, GraphPad Software, Boston, Massachusetts USA, www.graphpad.com. Statistical significance was defined as p < 0.05.

## RESULTS

### Overview of the HLA-DR3-specific PMT epitope analysis workflow

The first HLA class II associations with SjD were identified at the DR3 and DR2 loci in Caucasian populations, accounting for up to 90% of the MHC association. The HLA-DR3 allele, particularly the HLA-DRB1*0301-DQB1*0201 haplotype, has been consistently linked to SjD in various populations ^2–4,28–30^. Therefore, we focused on HLA-DR3 as we sought to determine the PMT associated with this specific HLA. To first define how post-translational modifications influence the landscape of HLA-DR3-presented epitopes derived from canonical SjD-associated autoantigens, we implemented a two-stage computational framework summarized in **Figure 1**. The analysis began with a reference set of 26 Ro60-, Ro52-, and La-derived peptides previously identified using the SMM approach as high-affinity binders to HLA-DR3^512^. This SMM-peptide dataset served as a defined baseline for evaluating PTM-dependent response. In addition, we utilized the full-length overlapping libraries for Ro60, Ro52, and La, covering 524, 460, and 394 amino acid antigens, respectively, to map modifiable anchor sites and PTM-sensitive regions across each antigen. The overlapping libraries were generated using the sliding-window approach, evaluated every position across Ro60, Ro52, and La, rather than restricting the analysis to annotated domains or a previously reported SMM-peptide dataset. In both datasets, peptides were screened for modifiable amino acids at canonical HLA-DR3 anchor positions (P1, P4, P6, and P9), expanded into PTM-mimic variants, and evaluated by comparing predicted binding affinities between native and modified sequences. Using both the reported epitope set and the full sliding-window scan yielded a unified, high-resolution map of PTM-sensitive positions.

**Figure 1.**
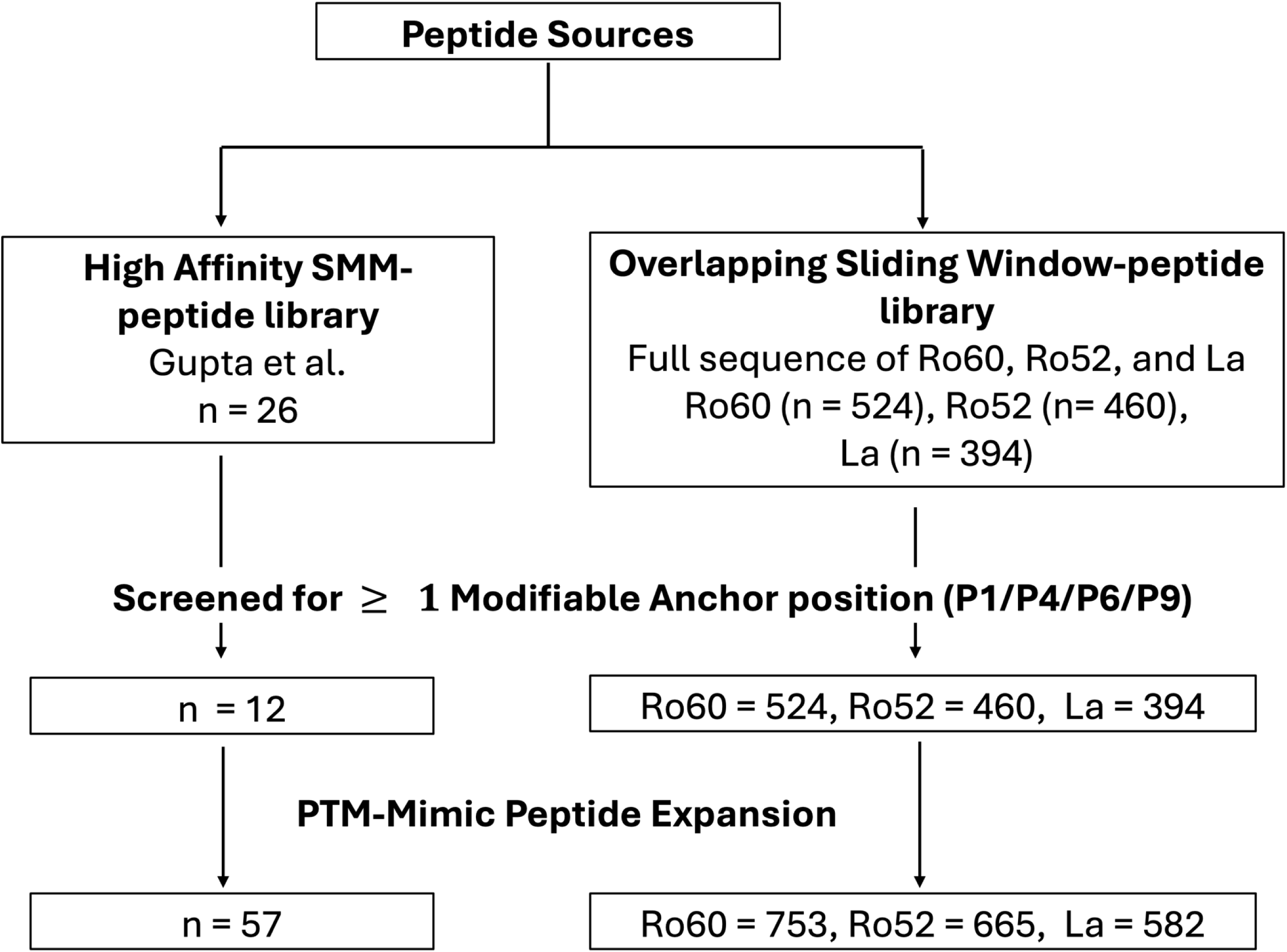
Overview of Peptide Libraries and PTM-Mimic Expansion Pipeline. Previously published high-affinity SMM-peptide library (Gupta et al.) and peptides obtained from an overlapping Sliding Window-peptide library covering the full-length Ro60, Ro52, and La sequence were combined to generate the initial peptide pool. The SMM-peptide dataset contains 26 peptides. The overlapping dataset contains 524 Ro60, 460 Ro52, and 394 La 15-mers, generated by sliding 1 amino acid at a time. All peptides were screened for the presence of at least one modifiable anchor residue at P1, P4, P6, or P9. The filtering step yielded 12 peptides from the SMM-peptide dataset, as well as 459 Ro60, 413 Ro52, and 347 La peptides from the overlapping dataset. Each retained peptide was subsequently expanded into its possible PTM-mimic variants, resulting in 57, 803, 717, and 656 mimic peptides from group 1, Ro60, Ro52, and La, respectively.

### Predictive binding affinity of PTM-mimic substitution in the SMM-peptide dataset

The SMM-peptide dataset comprises 26 previously reported HLA-DR3-restricted epitopes of Ro60, Ro52, and La autoantigens. All peptides contained at least one modifiable anchor residue and yielded a total of 56 PTM-mimic variants suitable for analysis. The native peptides showed a broad distribution of predicted affinities, with most values falling within a range typical for class II ligands of intermediate strength (**Figure 2, full data shown in Supplementary Table S1**). Substitution at PTM-eligible anchor residues rarely improved predicted binding. Instead, the majority of modified peptides exhibited higher IC₅₀ values than their corresponding native sequences. In total, 48 of 56 modified peptides (85.7%) showed negative log_2_(Native/PTM IC_50_) ratios, indicating reduced predicted affinity following substitution. The extent of these changes depended on both the identity of the residue and its anchor position. Mimics of arginine citrullination (R→Q) at P6 and P9 consistently resulted in weaker binding. Lysine acetylation mimics (K→Q) showed a similar pattern. Substitutions modeling phosphorylation (S/T→D) produced more modest effects, and oxidative mimics (F→Y, M→Q) typically resulted in small decreases that did not alter qualitative affinity class. Only a small subset of peptides showed any improvement, and these gains were limited in magnitude. Overall, PTM-mimic substitutions within the SMM-peptide epitopes produced limited changes to HLA-DR3 binding. Given the narrow dynamic range observed across this set, we next asked whether regions outside the annotated epitopes might harbor positions with stronger PTM sensitivity. This rationale motivated a comprehensive, sequence-wide analysis across the full Ro60, Ro52, and La proteins.

**Figure 2.**
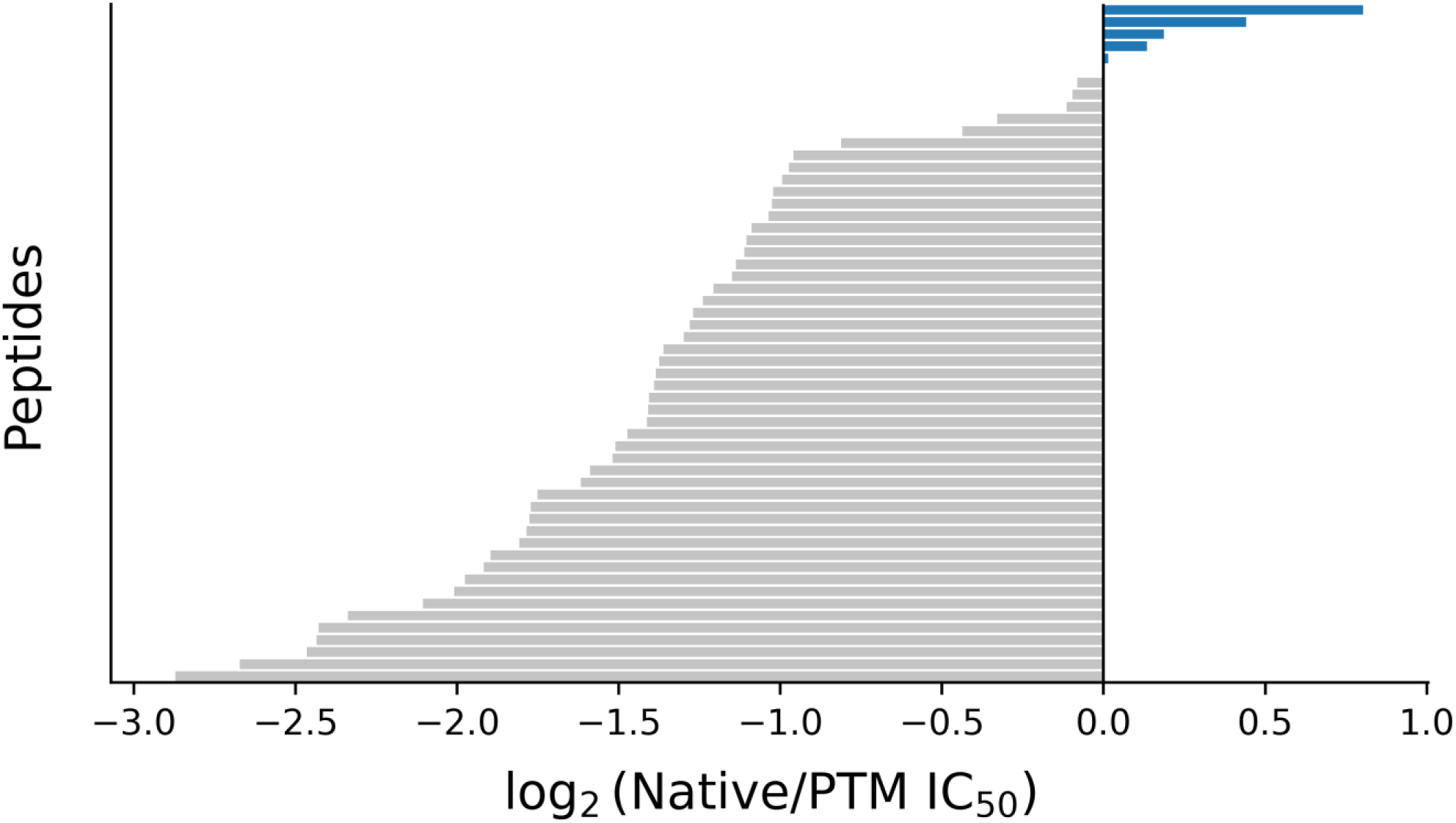
Predicted effects of PTMs on HLA-DR3 binding of the SMM-peptide epitopes. A waterfall plot shows the distribution of predicted changes in HLA-DR3 binding affinity for all PTM-mimic peptide variants derived from previously reported Ro60, Ro52, and La epitopes. Peptides are sorted by their log_2_ (Native/PTM IC_50_) values, where positive values (blue) indicate improved predicted binding after PTM substitution and negative values (grey) indicate reduced predicted binding. Each bar represents one WT-PTM peptide pair. A vertical line at log_2_ = 0 denotes unchanged predicted affinity.

### PTM-mimic substitution of the full-length Ro60, Ro52, and La autoantigens

After evaluating PTM-mimic substitution shifts within previously reported antigenic peptides, we applied the same anchor-based framework to full-length sequences to assess the frequency and distribution of PTM-eligible sites across the entire protein sequence. Extending the analysis to full-length sequences allowed us to systematically quantify the incidence of modifiable anchor sites of Ro60, Ro52, and La. Using the screening the sliding-window technique to generate the overlapping libraries identified 459 PTM-eligible peptides of Ro60, 413 PTM of Ro52, and 347 of La, defined as peptides containing at least one residue that could be substituted by a PTM-mimic at a canonical HLA-DR3 anchor position. We next summarized how these potential modifications were distributed across the four DR3 binding-motif positions (P1, P4, P6, and P9). To quantify the distribution of modifiable residues at these canonical anchor sites, we enumerated all possible substitutions at P1, P4, P6, and P9 across each overlapping library. The resulting positional counts are summarized in **Figure 3**. Across all three antigens, substitutions were not evenly distributed among anchor positions. P6 and P9 consistently exhibited the highest numbers of potential modifications, whereas P1 contributed the fewest. In Ro60, P6 and P9 accounted for 254 and 241 substitutions, respectively, compared with 169 at P4 and 139 at P1. Ro52 displayed a similar pattern, with 214 substitutions at P6 and 227 at P9, and lower counts at P4 (157) and P1 (119). La also showed elevated substitution counts at P6 (215) and P9 (167), relative to P4 (162) and P1 (111). These consistent patterns across Ro60, Ro52, and La establish the positional distribution of modifiable anchor residues prior to PTM-mimic expansion, indicating that potential PTM-permissive substitutions occur more frequently at P6 and P9 than at P1 or P4 across all three autoantigens.

**Figure 3.**
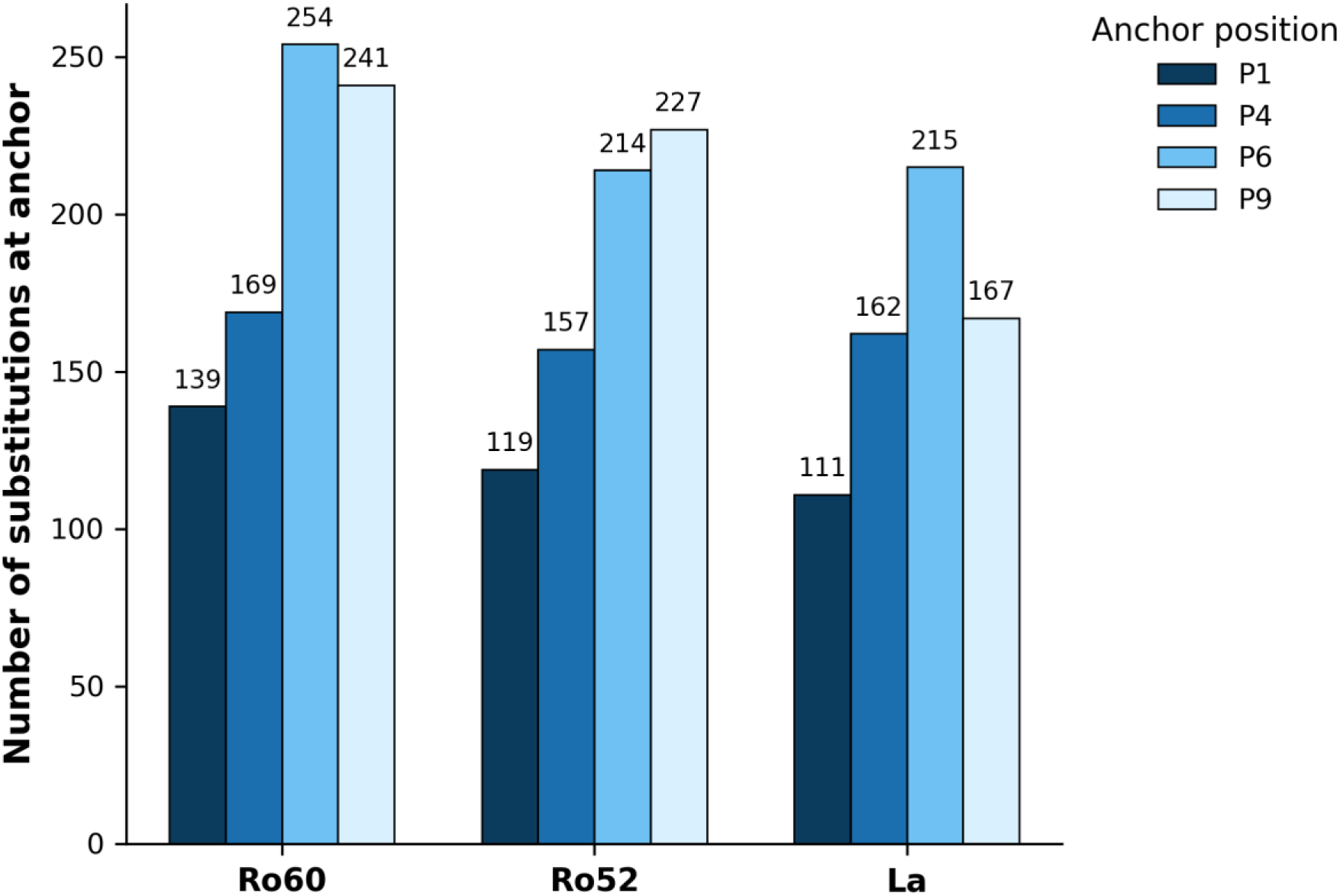
Distribution of PTM-mimic substitutions of the anchor positions. Counts of PTM-eligible substitutions at each HLA-DR3 anchor position (P1, P4, P6, and P9) across all sliding 15-mer peptides from Ro60, Ro52, and La. Bars indicate the number of sequence positions where an eligible residue occurs at the corresponding anchor site.

### Representation of PTM-mimic classes of Ro60, Ro52, and La autoantigens

To characterize the biochemical diversity of potential modifications, all PTM-eligible peptides were converted into mimic variants representing six predefined PTM classes: deamidation, phospho-mimic, acetylation-like, arginine-modification mimic, oxidation-like, and aromatic hydroxylation. This resulted in 803 PTM-mimic peptides for Ro60, 717 for Ro52, and 654 for La. As illustrated in **Figure 4A**, the representation of PTM classes varied considerably among the three antigens. Ro60 produced disproportionately large numbers of phospho-mimic (191) and acetylation-like (177) peptides, followed by arginine-modification mimics (124), aromatic hydroxylation (107), oxidation-like (104) peptides, and deamidation (100) peptides. Ro52 was enriched for deamidation-type (168) and phospho-mimic (143) variants, followed by arginine-modification mimics (133), whereas oxidation-like peptides were comparatively rare (40). In La, acetylation-like mimics predominated (212 variants), with smaller but still substantial contributions from deamidation (141) and phospho-mimic (124) variants. Oxidation-like (26) and arginine-modification mimics (63) occurred at much lower frequencies. These differences underscore that each antigen provides a unique substrate for PTM-driven substitution, shaped by its intrinsic residue composition and the distribution of anchor-accessible positions. The diversity of PTM classes represented also suggests that multiple biochemical pathways—including phosphorylation, deamidation, oxidation-like changes, and lysine modifications—could alter epitope presentation across these antigens.

**Figure 4.**
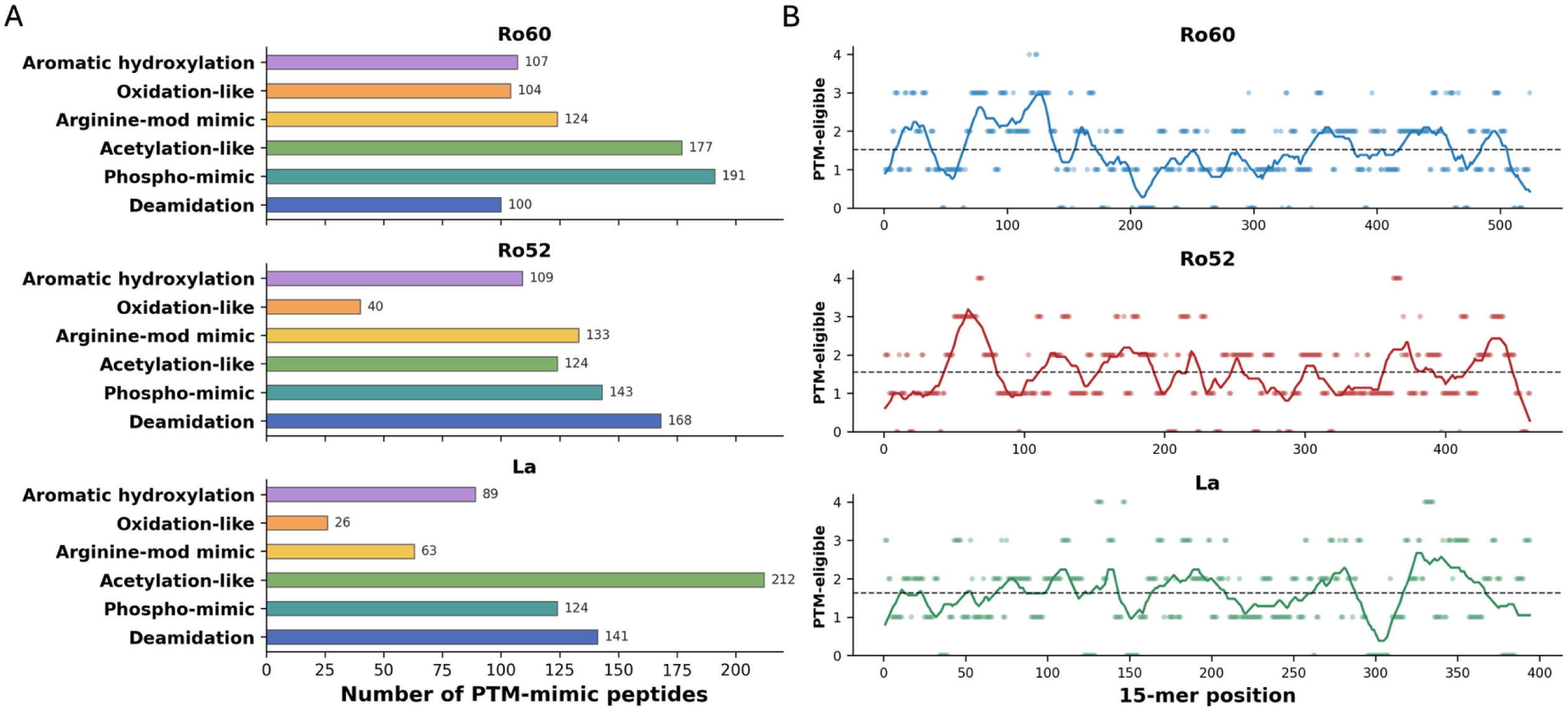
Categorical and positional distribution of PTM-mimic peptides. (**A**) Number of PTM-mimic peptides generated for each PTM class, shown separately for Ro60, Ro52, and La. PTM classes include aromatic hydroxylation, oxidation-like modifications, arginine-modification mimics, acetylation-like substitutions, phospho-mimics, and deamidation. (**B**) Distribution of PTM-eligible anchor sites along each protein sequence. Each scatter point corresponds to a single 15-mer peptide beginning at that sequence position and is plotted according to the number of DR3 anchor sites (P1, P4, P6, P9) within that peptide that contain PTM-eligible residues. Solid lines represent smoothed local averages, and black dashed lines indicate the mean number of PTM-eligible residues per protein.

We next examined how these PTM-eligible anchor sites were distributed across each antigen’s linear sequence. Mapping the number of modifiable anchor residues in each overlapping 15-mer revealed discrete high-density regions along all three antigens (**Figure 4B**). In Ro60, prominent peaks were centered approximately around residues 70-150, 355-390, 425-460, and 480-510. Ro52 exhibited multiple enriched intervals located near residues 50-90, 160-200, 250-285, and 420-455. La displayed three broader high-density regions, roughly spanning residues 70-150, 265-300, and 325-400. Windows containing three or more PTM-eligible anchors were largely confined to these intervals, whereas the remaining sequence showed lower and more uniform densities. These profiles indicate that PTM-eligible anchor sites cluster within defined sequence ranges rather than being evenly dispersed across each antigen (**Supplementary Table S2**).

### Domain-mapped PTM-eligible epitopes of Ro60, Ro52, and La autoantigens

To relate PTM-dependent changes in predicted HLA-DR3 binding to the structural organization of each antigen, we mapped log_2_(Native/PTM IC_50_) values onto annotated protein domains. The three antigens displayed distinct profiles, and each contained well-defined regions where PTM-mimic substitutions were associated with lower predicted IC_50_ values. For instance, as shown in **Figure 5A**, the strongest decreases in predicted IC_50_ of Ro60 were located within HEAT ring 2 and HEAT ring 3. These regions contained many windows with consistently negative log_2_(Native/PTM IC_50_) values across multiple PTM classes. The HEAT ring 1 and the distal C-terminal portion showed weaker and less consistent responses. The Y-RNA-binding interface also included several PTM-responsive windows, although the magnitude of these shifts was smaller. Ro52 displayed two major PTM-sensitive regions. One was located within the RING and B-box domains, and the other spanned the PRY-SPRY domain. Both regions contained groups of windows with lower predicted IC₅₀ values after PTM substitution. The coiled-coil segment between these domains showed only scattered responsive positions and produced smaller shifts (**Figure 5B**). Lastly, La showed a different distribution. Strong PTM-associated effects occurred within the La motif, RRM1, and parts of the central linker. These regions formed several clearly defined intervals with lower predicted IC_50_ values. Additional responsive positions were present in RRM2 and near the C terminus, but these changes were smaller and less continuous (**Figure 5C**). In summary, in the three antigens, positive log_2_(Native/PTM IC_50_) values were rare and appeared only at isolated positions. These domain-mapped profiles indicate that PTM-eligible epitopes are concentrated in specific structural regions rather than being evenly distributed along the antigen sequences.

**Figure 5.**
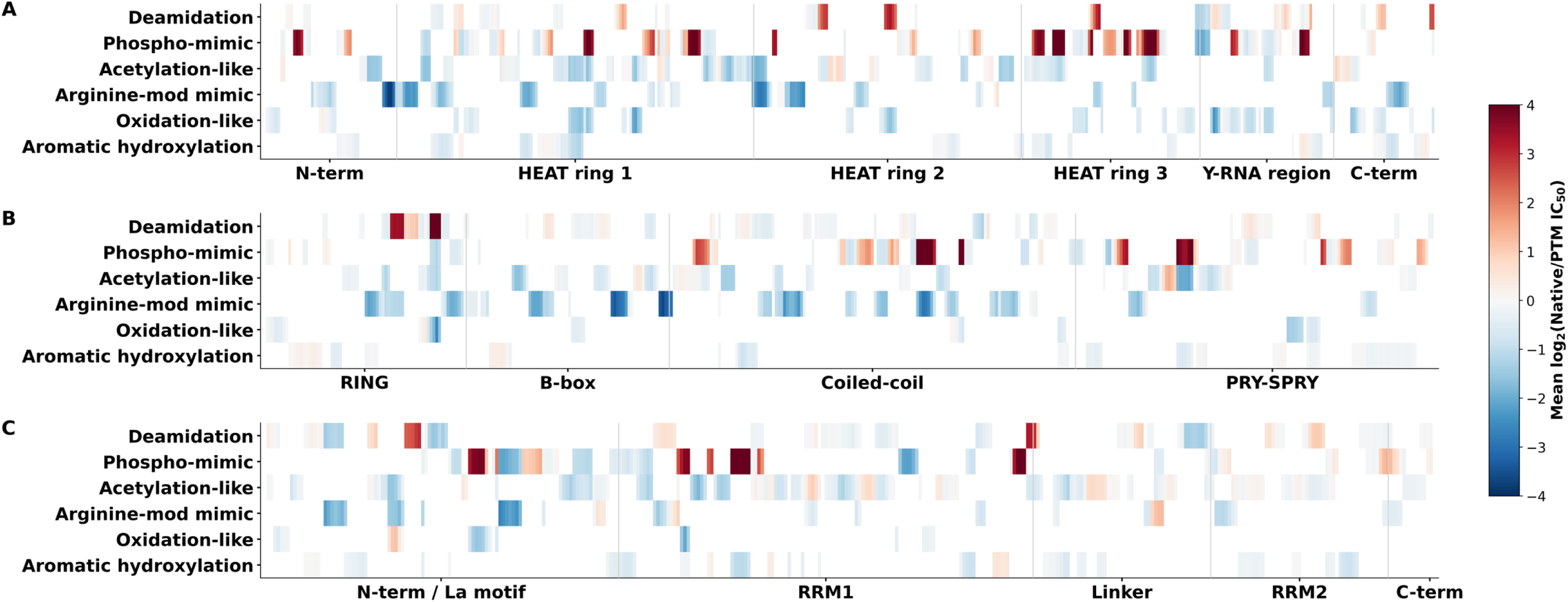
Domain-resolved landscape of PTM-eligible HLA-DR3-binding epitopes in Ro60, Ro52, and La autoantigens. Heatmaps depict the spatial organization of PTM-eligible regions across annotated domains of Ro60 (**A**), Ro52 (**B**), and La (**C**). Each tile represents the mean predicted change in HLA-DR3 binding affinity (log_2_ [Na/PTM IC_50_]) for PTM-mimic peptides whose modified anchor residues fall within the indicated sequence interval. Rows correspond to PTM classes (deamidation, phospho-mimic, acetylation-like, arginine-modification mimic, oxidation-like, and aromatic hydroxylation), and columns are aligned to protein domain boundaries. Positive values (red) indicate increased predicted binding following PTM substitution, whereas negative values (blue) indicate decreased predicted binding. Together, these maps reveal domain-associated patterns of PTM sensitivity within each antigen.

### Experimental validation of PTM-mimics showed enhanced T cell response

To experimentally assess whether predicted HLA-DR3 binding by PTM-mimics differences correspond to measurable T cell responses, and whether PTM-mimic substitutions capture functionally relevant features of the corresponding native PTMs, we selected four Native and PTM peptide pairs for experimental evaluation as a proof-of-concept. T cell responses were assessed using two complementary assays: IL-2 secretion as a measure of functional effector activity, and GFP reporter expression as a readout of TCR-dependent proliferation. The sequences and predicted HLA-DR3 binding affinities of the four Native-PTM peptide pairs are presented in **Table 1**. The first three pairs were derived from the overlapping analysis library of Ro60. These peptides were predicted to exhibit substantially increased HLA-DR3 binding affinity following PTM-mimic substitution compared with their corresponding native peptides. The last peptide pair was selected from the previously reported SMM-peptide dataset of Ro60 and was predicted to bind HLA-DR3 with similarly high affinity in both WT and PTM-mimic forms. As presented in **Figure 6A**, PTM-mimic peptides elicited higher IL-2 secretion than native peptides for pairs 1 and 2. For pair 1, IL-2 increased from 0.65 ± 0.39 pg/mL to 4.12 ± 0.52 pg/mL. In contrast, pairs 3 and 4 did not show increased IL-2 secretion in response to the native PTM peptides, with mean IL-2 decreased from 16.98 ± 1.31 to 13.37 ± 1.29 pg/mL for pair 3 and from 10.29 ± 3.01 to 6.35 ± 0.85 pg/mL for pair 4. The positive control induced robust IL-2 release, whereas blank and negative controls showed minimal activity. Using the GFP reporter assay to show proliferation, PTM-mimic peptides corresponding to pair 1 and 2 showed an increase in GFP^+^ events compared with native peptides. Pair 1 increased from 2281 ± 47 to 3361 ±253, and pair 2 increased from 1454 ± 184 to 3589 ± 322. For pairs 3 and 4, GFP responses were comparable between native and PTM-mimic peptides, indicating limited PTM-dependent effects on GFP^+^ activation under the conditions tested.

**Figure 6.**
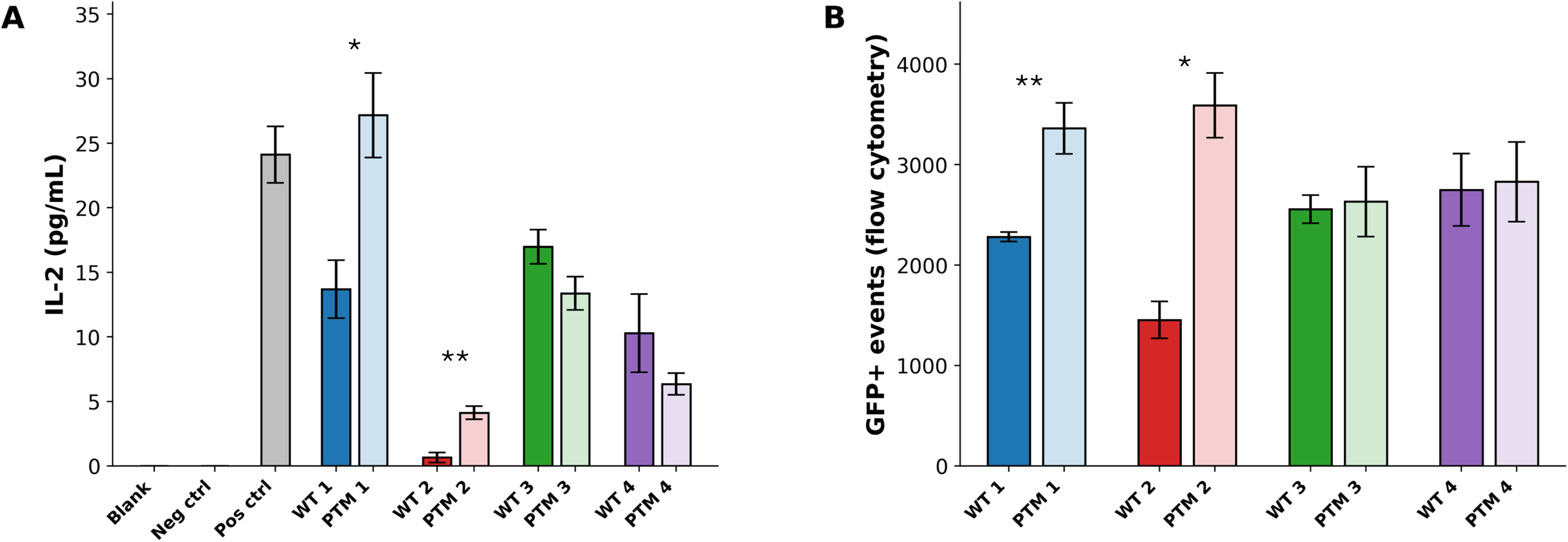
PTM-modified Ro60-derived peptides elicit functional and TCR-specific T cell responses. (**A**) Paired wild-type (WT) or native and PTM of Ro60-derived 15-mer peptides were compared for their ability to stimulate IL-2 production. WT (Native) and corresponding PTM peptides (peptides 1-4) were tested. The negative control was the ribonucleotide reductase subunit 1 from human alphaherpesvirus 1, spanning positions 709 to 721 (NVTWTLFDRDTSM). Blank samples had no peptides. Positive controls were a previously validated Ro60-derived 15-mer peptide with high predicted HLA-DR3 binding affinity. Bars represent mean ± SEM. (**B**) Disease-associated T cell receptors preferentially respond to PTM-modified Ro60-derived peptides. T cells expressing SjD-associated TCRs displayed enhanced activation in response to PTM peptides compared with matched Native peptides, measured as the number of GFP⁺ events by flow cytometry. Responses correspond to the same Native-PTM peptide pairs shown in panel (**A**). Bars represent mean ± SEM. Sequences and predicted HLA-DR3 binding affinities (IC_50_) of the WT (Native) and PTM peptides tested were provided in **Table 1**. *p≤0.05, **p≤0.01.

**Table 1.**
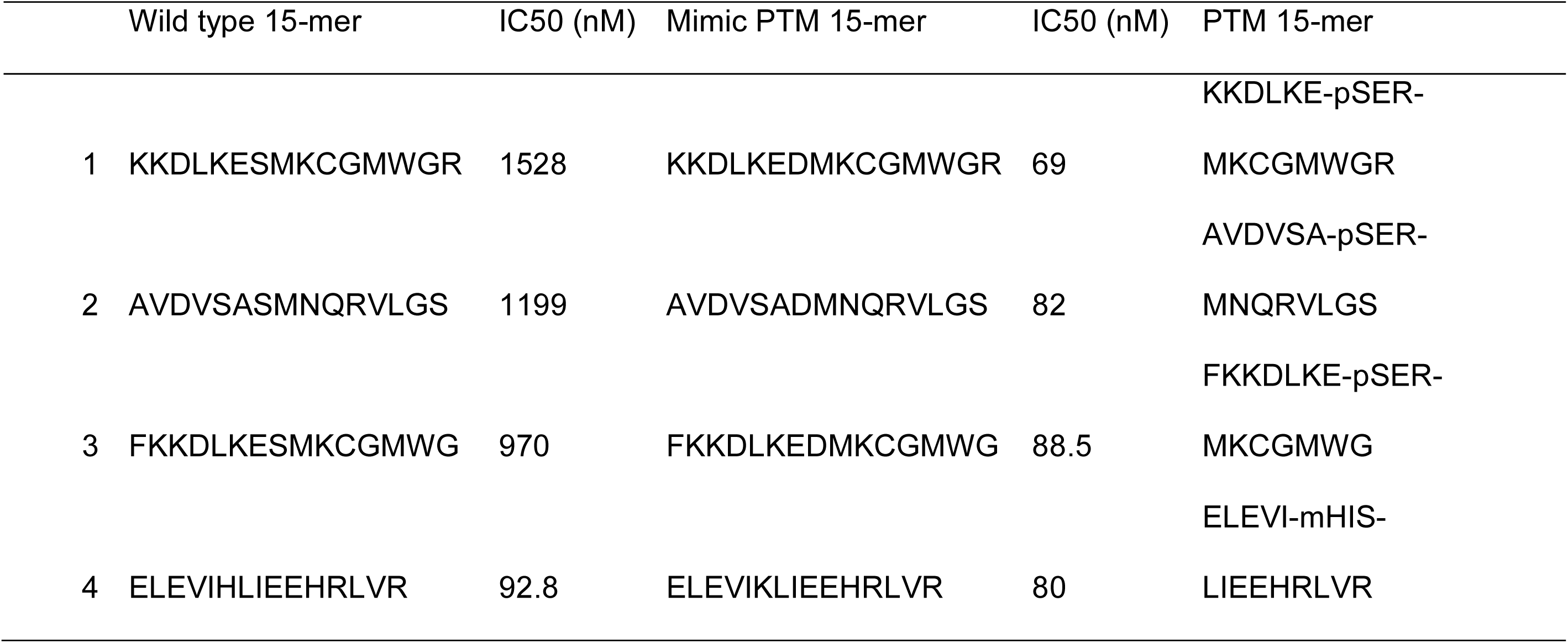
Peptide pairs selected for experimental validation and their predicted HLA-DR3 binding affinities.

### Structural modeling of a representative enhanced T cell response with PTM-mimic peptides

To investigate the structural basis for the enhanced T cell response, which is suggested by the better HLA-DR3 binding observed in the PTM-mimic peptide, we modeled the native and PTM-mimic 15-mers in complex with HLA-DR3 using the AlphaFold Server. Superposition of the two complexes showed that the overall peptide trajectory within the binding groove was preserved, but local deviations appeared around the substituted anchor residue (**Figure 7A**). The PTM-mimic peptide adopted a slightly shifted backbone orientation and altered side-chain positioning near the modified site. A close-up view of the modified region highlighted changes in local interactions (**Figure 7B**). The native anchor side chain was positioned close to a conserved pocket-lining residue in the HLA-DR3 binding groove (chain A Tyr11, ∼2.9 Å), whereas the PTM-mimic variant adopted a more distal orientation (∼6.7 Å). Despite this increased distance from PTM-mimic substitution, the PTM-mimic complex exhibits a larger overall peptide-DR3 interaction interface.

**Figure 7.**
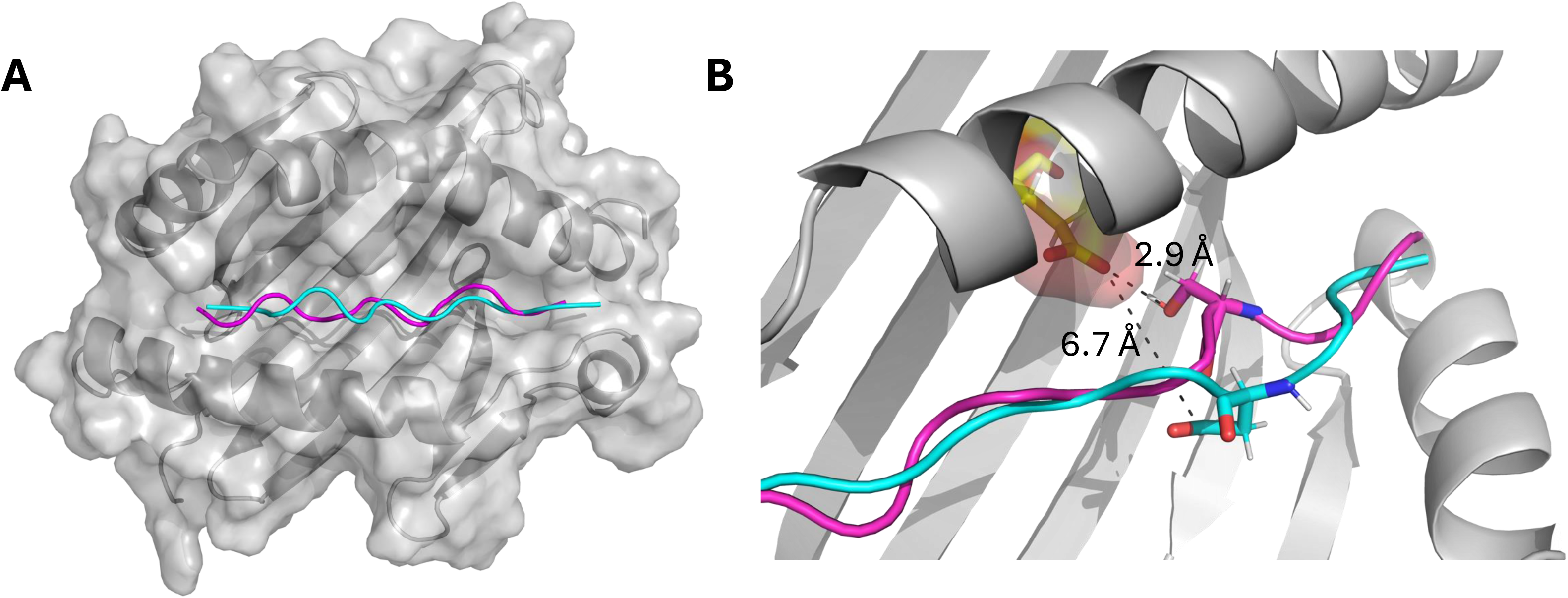
Structural comparison of WT and PTM-mimic peptides in the HLA-DR3 binding groove. (**A**) Superposition of the modeled Native (Native, magenta; KKDLKESMKCGMWGR, predicted IC_50_ = 1528 nM) and PTM-mimic (cyan; KKDLKEDMKCGMWGR, predicted IC_50_ = 69 nM) Ro60-derived 15-mer peptides bound to HLA-DR3. The two peptides adopt similar overall trajectories but diverge locally around the modified anchor residue, with the PTM-mimic showing a shifted register and altered side-chain orientation. (**B**) Enlarged view of the HLA-DR3 binding groove showing localized differences in anchor side-chain orientation between Native (magenta) and PTM-modified (cyan) peptides. The Native anchor side chain lies closer to a pocket-lining residue of the HLA-DR3 α chain (Tyr11), whereas the PTM variant adopts a more distal orientation. Distances are shown for reference. Measured distances are shown for reference to illustrate local differences in side-chain positioning.

Quantitative analysis using PDBSum revealed that the PTM mimics engage a greater number of interface residues and a larger contact surface with both HLA-DR3 chains (**Table 2**). Specifically, the peptide-DR3α interface area increased from 548 Å^2^ in the native complex to 688 Å^2^ in the PTM-mimic complex, accompanied by an increase in hydrogen bonds (6 to 9) and nonbonded contacts (109 to 132). Similarly, interactions with DR3β were expanded, with interface area increasing from 526 Å^2^ to 638 Å^2^, and nonbonded contacts from 70 to 96. Together, these changes reflect a broader engagement of the DR3 binding groove by the PTM-mimic peptides.

**Table 2.** Interface properties of WT and PTM-modified DR3-peptide complexes derived from PDBsum analysis.

| Model | Interface | Interface residues<br>(DR3 : peptide) | Interface<br>area (Å<sup>2</sup>) | Salt<br>bridges | Hydrogen<br>bonds | Non-bonded<br>contacts |
| --- | --- | --- | --- | --- | --- | --- |
| WT peptide:<br>KKDLKESMKCGMWGR | DR3α-<br>peptide |  |  |  |  |  |
|  | (A:C) | 17:11 | 548 | 0 | 6 | 109 |
|  | DR3β-<br>peptide |  |  |  |  |  |
|  | (B:C) | 15:7 | 526 | 3 | 7 | 70 |
| PTM peptide (S→D):<br>KKDLKEDMKCGMWGR | DR3α-<br>peptide |  |  |  |  |  |
|  | (A:C) | 20:14 | 688 | 2 | 9 | 132 |
|  | DR3β-<br>peptide |  |  |  |  |  |
|  | (B:C) | 18:09 | 638 | 4 | 8 | 96 |

## DISCUSSION

Post-translational modifications influence antigen processing and MHC class II presentation in several autoimmune conditions. Their contributions are most clearly defined in rheumatoid arthritis, where citrullinated and carbamylated peptides show enhanced binding to DRB1*04:01^4,31,32^. Studies with SLE samples have demonstrated similar effects with oxidized Ro60 and modified histones^5, 33^. Deamidated gliadin peptides in celiac disease provide another example of PTM-dependent improvements in MHC II affinity^34,35^. These findings established PTMs as modifiers of peptide binding and antigenicity, but their broader influence on the Ro60, Ro52, and La autoantigens of SjD has not been explored beyond isolated biochemical observations. Earlier studies focused on phosphorylation of Ro52^8^, oxidation of Ro60 (Scofield et al., 2005), or specific mimicry-related peptides^36^. None assessed how PTMs distributed across full antigen sequences might influence MHC binding, specifically the SjD-risk allele, HLA-DR3. This gap has limited the ability to evaluate whether PTM effects in SjD resemble patterns in other autoimmune diseases or whether Ro and La antigens have distinct features.

The current results address this gap by characterizing how PTMs shape predicted HLA-DR3 binding across the full lengths of Ro60, Ro52, and La. Among previously described peptides, PTM-associated affinity changes were common but modest, which is expected given that the SMM-peptide dataset consisted of high-affinity HLA-DR3 binders from Gupta et al^12^. High baseline affinity limits the dynamic range for further improvements. This behavior differs from celiac disease, where transglutaminase-mediated deamidation can convert weak HLA-DQ binders into strong binders^37^. In contrast, PTMs associated with rheumatoid arthritis and SLE tend to influence antigen processing, epitope exposure, or immune recognition rather than reliably increasing class II binding affinity^9,38^. The predominantly directional but modest shifts observed here align with prior analyses, which demonstrate that PTMs have limited effects when native affinity is already high^7^. These results established a rationale for a broader survey of the full Ro/La antigen sequences.

The sliding-window approach for overlapping peptide libraries revealed a diverse set of PTM-eligible anchor positions and more than 2,100 PTM-mimic peptides of the Ro60, Ro52, and La proteins. The predicted effects did not occur uniformly. Instead, they concentrated within structurally defined regions that differed among the three antigens. Ro60 displayed the strongest predicted responses within HEAT rings 2 and 3. Ro52 showed prominent clusters in the RING region, the B-box, and the PRY-SPRY domain. La exhibited enriched PTM-responsive intervals within the La motif, RRM1, and the central linker. These segments are characterized by flexibility, partial solvent exposure, or known roles in RNA or substrate recognition (Huang et al., 2010). Similar associations between PTM responsiveness and structural dynamics have been reported for modified autoantigens in SLE and rheumatoid arthritis (Utz et al., 2000; Doyle and Mamula, 2012; Tilvawala et al., 2018). These parallels suggest that PTMs in SjD are likewise most consequential in regions naturally accessible to modification and antigen processing.

These structural properties align closely with principles of MHC class II antigen presentation. MHC class II immunodominance requires that peptides arise from segments that are accessible, protease-sensitive, and capable of adopting registers compatible with the DR3 groove (Neefjes et al., 2011). Each of the PTM-responsive regions identified above fulfills these criteria: the Ro60 HEAT-repeat solenoid undergoes small conformational shifts during RNA engagement; the PRY-SPRY domain of Ro52 contains surface loops that readily adjust; and the La motif and RRM1 include partially exposed regions that reposition during RNA binding^37–39^. PTMs arising in these structurally permissive environments may therefore stabilize favorable registers or enhance compatibility with pockets P1, P4, P6, or P9. This mechanism mirrors findings from other autoimmune conditions in which PTMs strengthen pocket interactions or reposition side chains to promote class II loading^40^

Experimental validation of four representative Native-PTM pairs largely supported the computational predictions of PTM-dependent response of HLA-DR3 binding. For three of the four pairs tested (Pair 1, 2, and 4), PTM-mimic peptides consistently exhibited higher experimental activity than their corresponding native peptides, in agreement with the predicted increases in HLA-DR3 binding affinity. This observation was supported by the GFP proliferation assay, which directly reports TCR-dependent activation and showed distinction between native and PTM peptides. However, pair 3 deviated from this overall pattern. Although the PTM-mimic peptide was predicted to exhibit enhanced HLA-DR3 binding, the experimental observation difference between native and PTM peptides was reduced. Together, these results support a PTM-mimic-based screening strategy for identifying peptides associated with PTMs that exhibit enhanced T cell activity. The experimental data demonstrate that peptides identified by this screening could exhibit measurable functional differences when tested experimentally, consistent with predicted PTM-dependent effects on HLA-DR3 binding.

To explore whether the experimental PTM-dependent effect on HLA-DR3 binding and T cell activation could be rationalized at a structural level, we performed comparative modeling of one representative Ro60 native-PTM peptide pair binding to HLA-DR3. Although the PTM-modified side chain is positioned further from the conserved pocket-lining residue (chain A Tyr 11) in the HLA-DR3 binding groove, quantitative interface analysis indicates that this local change is accompanied by an overall expansion of peptide-HLA-DR3 contact. In the modeled PTM-mimic complex, the increased contact area and more hydrogen and non-hydrogen bonds tend to form a more stable interaction between the peptide and both HLA-DR3 chains. These observations suggest that PTM-associated alterations at individual anchor positions can be accommodated through redistribution of contacts along the binding groove, rather than through a single dominant interaction. Importantly, this structural example is not intended to define a general mechanism but instead illustrates how specific PTMs can enhance peptide compatibility with the HLA-DR3 groove through the modest, localized adjustment. These adjustments did not involve major changes in backbone geometry, which is consistent with established structural determinants of class II peptide binding^40^. When considered in conjunction with experimental validation, the structural analysis supports the interpretation that PTM-mimic-based screening can identify peptides whose native post-translational modifications promote productive engagement with HLA-DR3.

Several limitations should be noted. PTM-mimic substitutions approximate but cannot fully reproduce native chemical modifications. Only four peptide pairs were experimentally validated, and the functional consequences for antigen presentation and T-cell activation were not evaluated. Structural modeling was performed for a single pair and is presented illustratively rather than comprehensively. Future studies involving antigen-presentation assays, T-cell functional analyses, and in vivo PTM mapping in salivary gland tissue will be necessary to determine which modified peptides are processed and recognized during disease development.

Together, these observations connect SjD to PTM-dependent changes in antigen presentation seen in other autoimmune diseases. Salivary gland epithelial cells in SjD experience oxidative stress, disrupted phosphorylation pathways, and lipid-reactive oxygen species accumulation (Zhou et al., 2024). These conditions favor multiple forms of PTM, and the correspondence between PTM-responsive regions identified here and structurally accessible domains suggests that modified peptides could be generated in vivo. Such peptides may contribute to the heterogeneous CD4⁺ T-cell responses reported in the disease and are consistent with earlier evidence of epitope diversification in Ro and La autoreactivity^9^. Although additional functional studies are required, these findings provide conceptual support for considering PTM - modified Ro and La peptides as potential contributors to loss of tolerance in SjD. In conclusion, this work characterizes how PTMs influence the HLA-DR3-restricted epitope landscape of Ro60, Ro52 and La. The integration of sequence-wide scanning, PTM-mimic analysis, experimental testing, and structural modeling identifies discrete PTM-eligible regions of the SjD-associated autoantigens. These findings suggest that PTMs can alter the repertoire of peptides available for class II presentation in SjD, providing a foundation for future work aimed at identifying PTM-dependent neoepitopes relevant to disease pathogenesis.

## AUTHOR CONTRIBUTIONS

DL and CN conceived the study and designed the experiments. DL designed and performed the computational studies and analysis. AV performed the in vitro experimental validation experiments. All authors reviewed and approved the manuscript.

## ACKNOWLEDGMENTS

Research reported in this publication was supported by the National Institute of Dental & Craniofacial Research of the National Institutes of Health under Award Numbers R56DE028544 and R01DE028544. The content is solely the responsibility of the authors and does not necessarily represent the official views of the National Institutes of Health.

